# Nucleolar targeting in an early-branching eukaryote suggests a general physicochemical mechanism for ribosome protein sorting

**DOI:** 10.1101/2021.12.20.473284

**Authors:** Milad Jeilani, Karen Billington, Jack Daniel Sunter, Samuel Dean, Richard John Wheeler

**Affiliations:** Sir William Dunn School of Pathology, University of Oxford, OX1 3RE; Tunbridge Wells Hospital, Tonbridge Rd, Tunbridge Wells, TN2 4QJ; Department of Biological and Medical Sciences, Oxford Brookes University, OX3 0BP; Warwick Medical School, Warwick University, CV4 7AL; Peter Medawar Building for Pathogen Research, Nuffield Department of Medicine, University of Oxford, OX1 3SY

**Keywords:** nucleolus, nucleolar targeting, *Trypanosome*, mitochondrial ribosome, liquid-liquid phase separation

## Abstract

The eukaryotic cell targets proteins to the organelles in which they function, both membrane-bound (like the nucleus) and non-membrane-bound (like the nucleolus). Nucleolar targeting relies on positively charged localisation signals, and has received rejuvenated interest since the widespread recognition of liquid-liquid phase separation (LLPS) as a mechanism contributing to nucleolus formation. Here, we exploit a new genome-wide analysis of protein localisation in an early-branching eukaryote, *Trypanosoma brucei*, to analyse general nucleolar protein properties. *T. brucei* nucleolar proteins have similar properties to those in common model eukaryotes, specifically basic amino acids. Using protein truncations and addition of candidate targeting sequences to proteins, we show both homopolymer runs and distributed basic amino acids give nucleolar partition, further aided by a nuclear localisation signal (NLS). These findings are consistent with phase separation models of nucleolar formation and protein physical properties being a major contributing mechanism for eukaryotic nucleolar targeting, conserved from the last eukaryotic common ancestor. Importantly, cytoplasmic ribosome proteins in comparison to mitochondrial ribosome proteins followed the same pattern – pointing to adaptation of physicochemical properties to assist segregation.

**Summary Statement:** We show protein targeting to the nucleolus is mediated by positive charge, likely across eukaryotes, and contributes to sorting of mitochondrial from cytoplasmic ribosome proteins.

## Introduction

The nucleolus or nucleoli are typically the largest non-membrane bound compartments within the nucleus, a site for ribosome biogenesis, and are found near-universally in eukaryotes. The unicellular parasite *Trypanosoma brucei* is no exception. This early-branching eukaryote causes African trypanosomiasis (sleeping sickness) in humans and nagana in animals. A prerequisite for specialised function of any organelle is protein partitioning, but protein features defining targeting to the nucleolus and whether they are conserved in early-branching eukaryotes are incompletely understood.

Nucleolar targeting is of particular interest in *T. brucei* as they have life cycle stage-associated adaptation of nucleolar structure. Unusually, *T. brucei* uses RNA polymerase I (Pol I) and basal Pol I transcription factors for transcription of the major surface antigen protein-coding gene(Günzl et al., 2003; Pays et al., 1989) in addition to ribosomal RNA (rRNA) precursors. In some life cycle stages, this is closely associated with the nucleolus (procyclic form, procyclin), while in others it is in a dedicated nuclear body distinct from the nucleolus (bloodstream form, variant surface glycoprotein)(Daniels et al., 2010; Landeira and Navarro, 2007; Navarro and Gull, 2001a). *T. brucei* can therefore control whether certain proteins partition to only the nucleolus, or also a non-nucleolar compartment with additional specific components.

Many nuclear localisation signals (NLSs) are a short linear motif(Dingwall et al., 1982; Mattaj and Englmeier, 1998), but a similar nucleolar localisation signal (NoLS) has remained elusive. Many sequences are known to be NoLSs, typically identified through motif deletion mutants(Duan et al., 2019; Iyama et al., 2018; Musinova et al., 2011; Savada and Bonham-Smith, 2013; Schmidt-Zachmann and Nigg, 1993), and are used as the basis of predictive tools(Scott et al., 2010; Scott et al., 2011). Generally, NoLSs are positively charged (many basic amino acids), and it is proposed that electrostatic interactions confer nucleolar enrichment(Musinova et al., 2011; Musinova et al., 2015; Savada and Bonham-Smith, 2013). However, high NoLS sequence diversity and distinguishing NLSs and NoLSs (both have many basic amino acids) make NoLS prediction challenging(Martin et al., 2015).

The nucleolus has liquid-like properties(Brangwynne et al., 2009; Brangwynne et al., 2011; Feric et al., 2016), and recent work has shown that nucleolar proteins can undergo liquid-liquid phase separation (LLPS) in vitro(Boeynaems et al., 2018; Feng et al., 2019; Gomes and Shorter, 2019; Lafontaine et al., 2021; Sawyer et al., 2019b; Wang and Zhang, 2019; Woodruff et al., 2018). LLPS as a model for nucleolus formation provides a new conceptual framework for understanding nucleolar targeting(Feric et al., 2016; Lafontaine et al., 2021). In the LLPS model, key abundant components termed ‘scaffolds’ have characteristic physicochemical properties which lead to their phase separation under cellular conditions, giving a condensate phase with an up to 100-fold higher concentration of scaffold(Li et al., 2012; Nott et al., 2015). Building on this model, the condensate is a different environment to the surrounding cytoplasm, defined by the physicochemical properties of the scaffold, and favourable interaction or solvation in the condensate allows partition of ‘client’ proteins into the condensate(Feng et al., 2019; Lafontaine et al., 2021; Martin and Holehouse, 2020).

Multivalent scaffold-scaffold interaction is important for LLPS and often arises from intrinsically disordered regions (IDRs)(Banani et al., 2017; Boeynaems et al., 2018; Duan et al., 2019; Holehouse et al., 2017; Lin et al., 2018; Stenström et al., 2020). IDRs are over-represented in membraneless organelle proteins, including nucleolar proteins, particularly those able to undergo phase separation in vitro(Lin et al., 2015; Martin and Mittag, 2018; Meng et al., 2015; Sawyer et al., 2019a; Stenström et al., 2020; Wright and Dyson, 2015). IDR amino acid composition tends to be better conserved than primary sequence suggesting physicochemical properties rather than simply electrostatic interactions dictate LLPS behaviour(Martin and Mittag, 2018; Weber, 2017). Growing evidence that nucleolar protein IDRs drive partition to the nucleolus phase(Duan et al., 2019; Stenström et al., 2020) may explain the lack of a specific NoLS sequence.

We suggest an early-branching eukaryote like *T. brucei* will give more insight to protein partition to the nucleolus, in addition to the importance of a species-specific model. Previous analysis of NLSs in *T. brucei* have convincingly shown the canonical monopartite NLS (K-^K^/_R_ -X-^K^/_R_) in model eukaryotes(Chelsky et al., 1989) is functional in *T. brucei*(Marchetti et al., 2000) and classical NLSs are strongly enriched in *T. brucei* nuclear proteins identified by mass spectrometry(Canela-Pérez et al., 2019; Goos et al., 2017). *T. brucei* and *T. cruzi* α-importin has also been shown to bind to a bipartite NLS(Afrin et al., 2020; Canela-Pérez et al., 2018; Canela-Pérez et al., 2020). NLS conservation in such an early-branching eukaryote strongly suggests that this is the ancestral nuclear transport mechanism, and likely common across eukaryotes. We have applied similar logic to understand NoLSs.

Here, we exploit genome-wide localisation data from our *T. brucei* localisation database, TrypTag(Dean et al., 2017), to quantify enrichment of a tagged copy of every *T. brucei* protein in the nucleus and nucleolus. Re-identification of the canonical NLS validated this approach, and it identified basic amino acids as the key protein feature associated with nucleolar partition, both in short IDRs and distributed through a protein, and importantly in an early-branching eukaryote. This suggests protein charge is the mechanism for nucleolar targeting across all eukaryotes, consistent with a LLPS model of nucleolar formation forming an environment which promotes partition of basic client proteins. Importantly, mitochondrial ribosome (mitoribosome) and cytoplasmic ribosome (cytoribosome) proteins also follow this pattern suggesting a contributing mechanism to their localisation.

## Results

### A genome-wide map of protein partition into the nucleus and nucleolus

The TrypTag genome-wide protein localisation project(Dean et al., 2017) has generated tagged cell lines and captured high resolution microscope images of cell lines expressing endogenously tagged copies of 89% of *T. brucei* proteins (excluding variant surface glycoproteins). Protein tagging with mNeonGreen (mNG) was attempted at both the N and C terminus, with N and C terminal data available for >75% of cell lines. Each cell line was recorded through diffraction limited widefield epifluorescence images of multiple fields of view, typically ≥4 fields of view typically containing ≥250 cells – ∼5 million cells in total.

We use automated high content image analysis to analyse the partition of proteins to the nucleus vs. cytoplasm and partition to the nucleolus vs. nucleoplasm. The nucleus was identified using signal from the DNA stain Hoechst 33342, and the nucleolus center identified from the darkest point near the centre of the nucleus. Per cell, sum signal intensity from mNG fluorescence was calculated for the cytoplasm and nucleus, with nuclear signal further broken down to sum nucleoplasm or nucleolar signal intensity. Each cell was analysed individually then averaged to generate per-cell line (ie. per N or C terminally tagged protein) data.

To validate the quality of the data we plotted total mNG signal in the cell against nucleus/cytoplasm partition and nucleolus/nucleoplasm partition (Figure 1A). Distinct populations with high nucleus/cytoplasm and high nucleolus/nucleoplasm partition were readily visible and manual validation confirmed that these were nuclear and nucleolar proteins respectively (Figure 1B,C). A weak positive correlation of nucleus/cytoplasm partition for nuclear proteins was visible – likely arising from a constant cell autofluorescence background.

**Figure 1.**
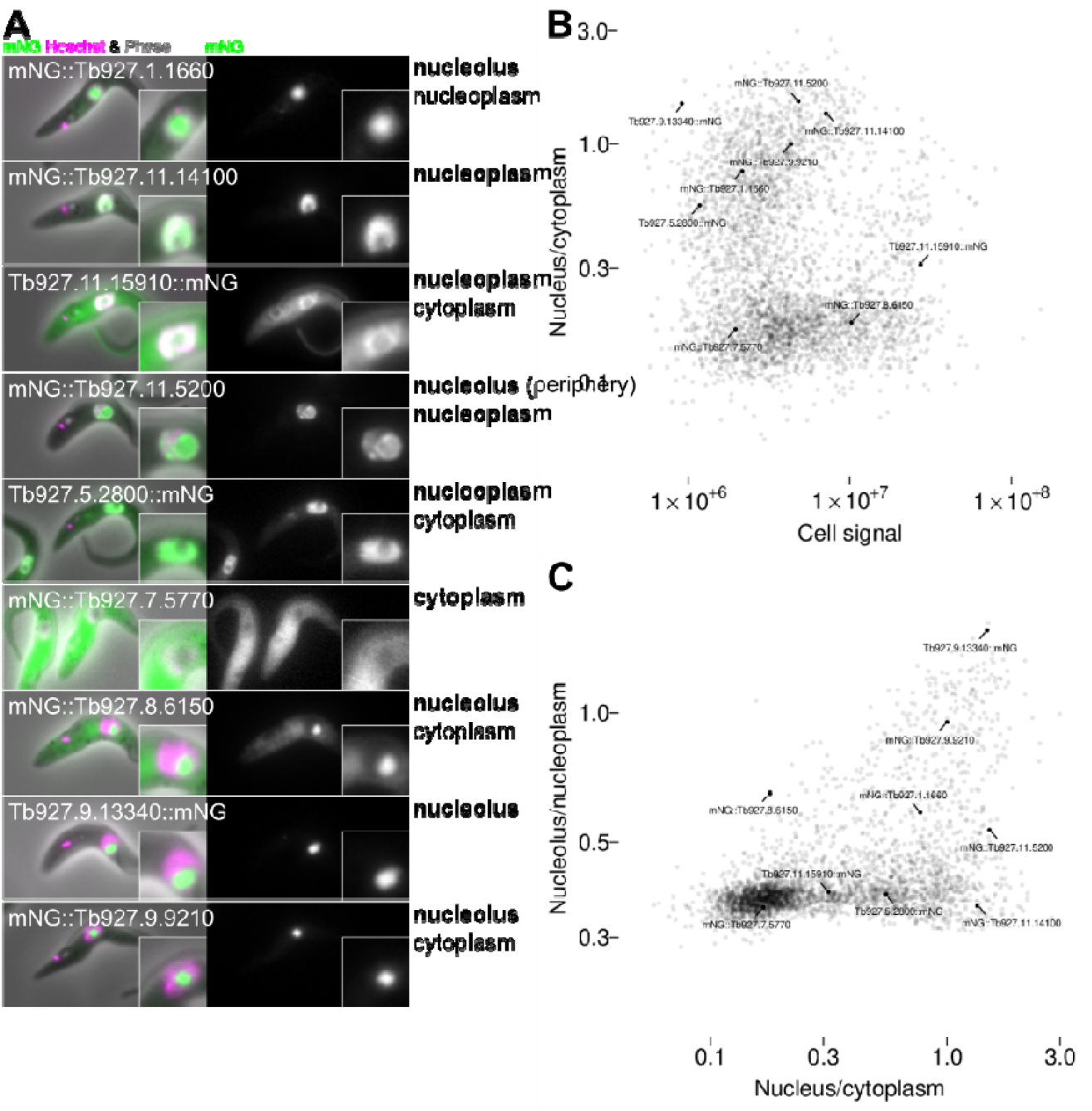
The appearance of nucleolar, nucleoplasmic and cytoplasmic proteins in *T. brucei*. A. mNG tagged *T. brucei* proteins showing a range of proteins with nucleolar, nucleoplasmic and/or cytoplasmic proteins. B. Cell signal and nucleus/cytoplasm partition from Figure 2A replotted showing the cell lines in A. C. Nucleus/cytoplasm and nucleolus/nucleolus partition from Figure 2C replotted showing the cell lines in A.

The nuclear pore is a diffusion barrier, expected to reduce nuclear access for proteins >60 kDa. The mNG tag and linker is ∼30 kDa, therefore we may see a threshold around ∼30 kDa untagged molecular weight in nuclear/cytoplasm partition behaviour. However, we saw no clear correlation of nuclear/cytoplasm partition with molecular weight (Figure 2B).

**Figure 2.**
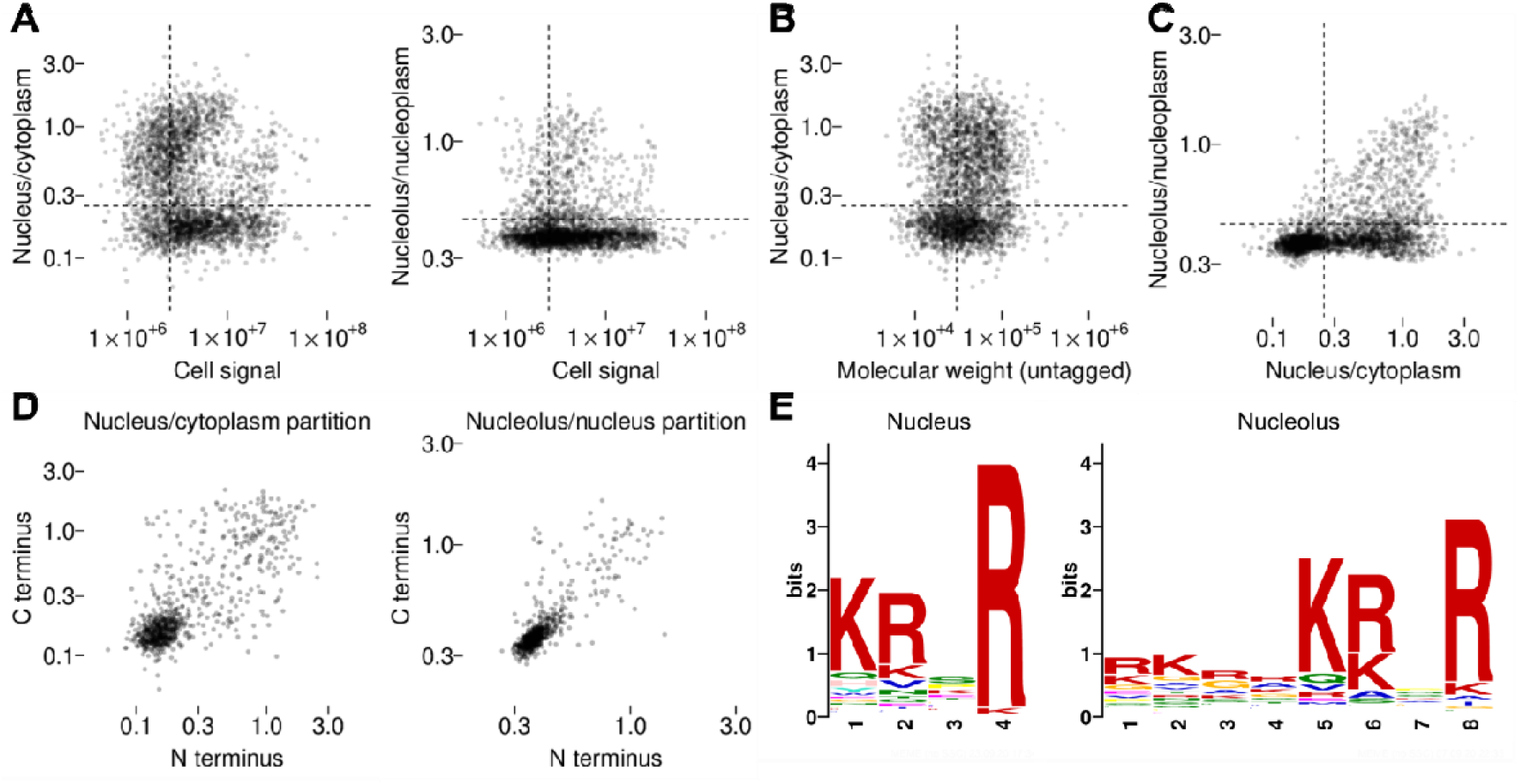
Protein motifs in nucleus and nucleolus *T. brucei* proteins identified from automated protein localisation quantitation. A. Automated quantitation the ratio of nucleus to cytoplasm mNG fluorescence signal or the ratio of nucleolar to nucleoplasm mNG fluorescence plotted against sum mNG fluorescence signal. The average signal and nucleus/cytoplasm ratio for all cells (typically >200 cells) for all cell lines in the TrypTag protein localisation database manually annotated as localising to the nucleolus, nucleoplasm and/or cytoplasm (>3000 cell lines) is plotted. Nucleus/cytoplasm ratio or nucleolus/nucleoplasm ratio was used to classify proteins as nuclear and/or nucleolar respectively, the dashed line represents the cutoff. B. Nucleus/cytoplasm ratio plotted against protein molecular weight, ignoring the mNG tag molecular weight. There is no clear correlation between molecular weight and nucleus/cytoplasm partition. The dashed lines represent the nucleus/cytoplasm nuclear classification cutoff and an approximate cutoff for protein expected to be too large to diffuse through the nuclear pore. C. Nucleolus/nucleoplasm ratio plotted against nuclear/cytoplasm ratio. The dashed lines represent the nucleus/cytoplasm nuclear classification and nucleolus/nucleoplasm nucleolus classification cutoffs. D. Correlation of nucleus/cytoplasm ratio and nucleolus/nucleoplasm ratio for all cell lines in the TrypTag database where the same protein tagged at either the N or C terminus with mNG gave signal intensity above the background intensity (>700 cell line pairs). E. Protein motifs identified by MEME(Bailey et al., 2009) for proteins above the nucleus/cytoplasm and nucleolus/nucleoplasm cutoffs.

For ongoing analysis, we defined nucleus/cytoplasm and nucleolus/nucleoplasm partition thresholds to classify proteins as nucleolar, nucleoplasmic, nuclear (nucleolar or nucleoplasmic) or cytoplasmic (neither nucleolar nor nucleoplasmic) (Figure 2C). These were selected as inclusive thresholds, e.g. a tagged protein was classified as nucleolar so long as it has high nucleolar/nucleoplasm partition, but may also have easily visible nucleoplasmic and/or cytoplasmic signal.

For many proteins, both the N and C terminally tagged cell lines were successfully generated. We compared the nucleus/cytoplasm and nucleolus/nucleoplasm partition for N and C terminally-tagged proteins, which showed a good positive correlation albeit with some outliers (Figure 2D). For ongoing analysis, we treated evidence from N or C terminal tagging independently – essentially classifying a protein as nucleolar or nucleoplasmic if there was evidence from either terminus.

Using these nucleolar and nucleoplasmic gene lists we searched for motifs enriched in each set using MEME(Bailey et al., 2009). This identified one statistically significant motif for each list – the canonical nuclear localisation signal (NLS): KRXR.

### KRXR is necessary and sufficient for protein targeting to the nucleus

To determine whether KRXR is necessary for protein targeting to the nucleus, we searched the genome for proteins with this motif near (contained within 15 amino acids of) either the N or C terminus. We selected these proteins as it is possible to remove the candidate NLS through a small open reading frame (ORF) truncation at the endogenous locus using PCR based approach.

We selected 8 genes with a single candidate NLS near the N or C terminus. Truncation of these genes to remove the NLS (and introduce an mNG tag) is a test of whether the candidate NLS is sufficient for nuclear localisation (Figure 3A). All except one protein (mNG::Δ9-Tb927.3.1350) showed the NLS was necessary for a strong nuclear localisation, although only two proteins (mNG::Δ13-Tb927.10.12980 and mNG::Δ11-Tb927.10.3970) appeared excluded from the nucleus in the absence of their NLS. We selected a further 3 genes with multiple candidate NLSs where only one candidate NLS is near the N or C terminus (Figure 3B). We expect these N or C terminal NLSs not to be necessary for the proteins’ nuclear localisation, and indeed all were not necessary for a strong nuclear localisation.

**Figure 3.**
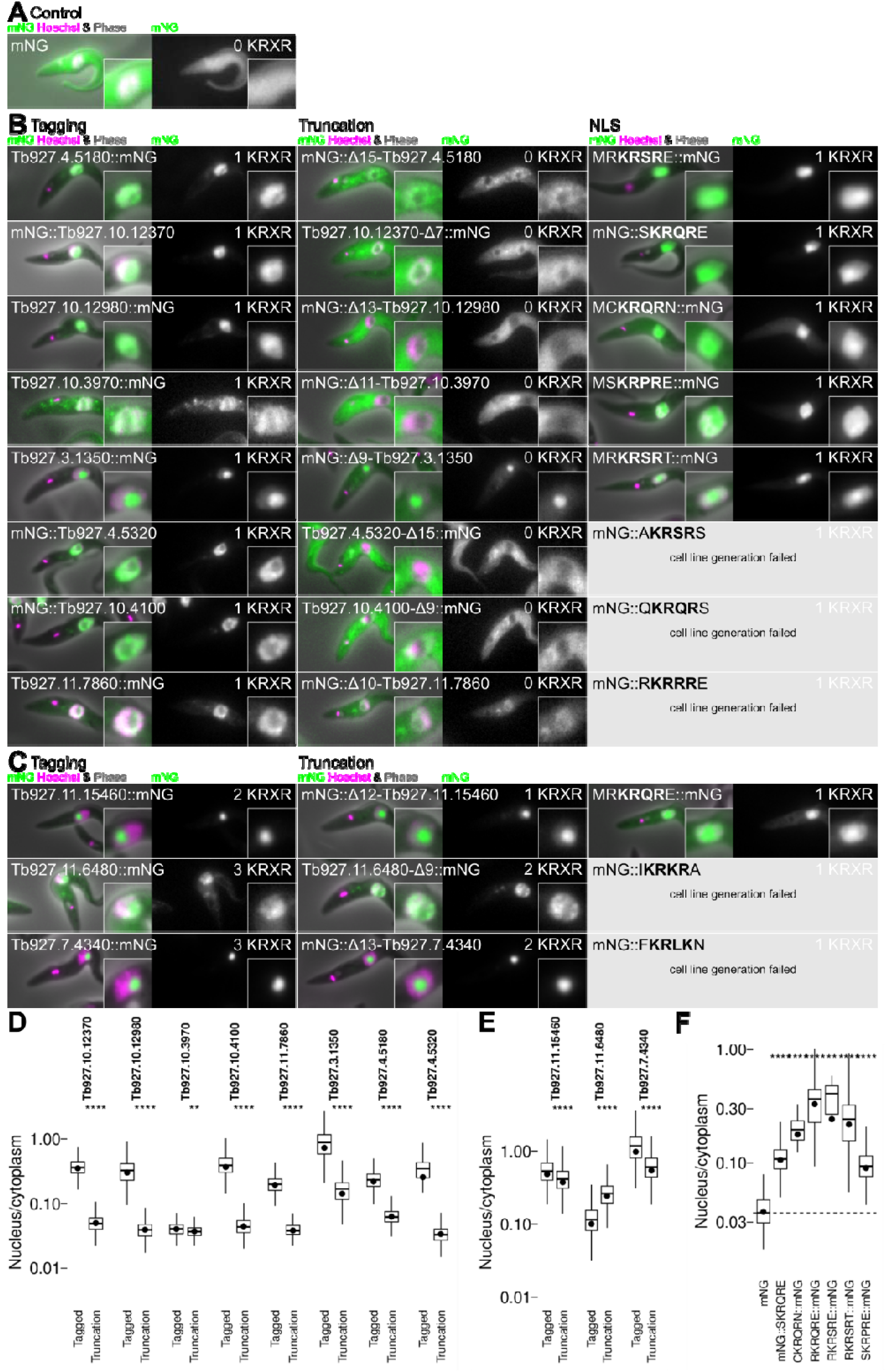
The canonical KRXR NLS is a functional NLS sufficient and necessary for targeting of many proteins to the nucleus. A. Localisation of mNG when expressed in *T. brucei* cells. B. Testing the candidate NLS found in 8 nuclear proteins with a single candidate NLS near the N or C terminus. Localisation of the protein by tagging at the endogenous locus; localisation following truncation to remove the NLS and replacement with mNG; and localisation of mNG fused to the candidate NLS. For each cell line, the number of KRXR motifs in the mNG fusion is shown in the top right. C. As for B, except for 3 nuclear proteins with multiple candidate NLSs, one of which is near the N or C terminus. D. Plots of automated quantitation of the nucleus/cytoplasm mNG fluorescence signal partition from the cell lines in B. E. Plots of automated quantitation of the nucleus/cytoplasm mNG fluorescence signal partition from the cell lines in C. F. Plots of automated quantitation of the nucleus/cytoplasm mNG fluorescence signal partition from the cell line in A, and those successful cell lines in B and C that involve mNG fused to a candidate NLS Statistical significance was assessed using the Wilcoxon signed-rank test (ns not significant, * p≤0.05, ** p≤0.01, *** p<0.001, **** p≤0.0001).

To test whether these NLSs are sufficient to confer a nuclear localisation, we generated cell lines expressing mNG fused to the NLS from each gene. In each case, we took the NLS sequence with one flanking amino acid and fused it to the mNG N or C terminus based on where it was found in the source gene. By this approach, we had a low success rate generating cell lines with an NLS fused to the mNG N terminus. However, all successfully tested NLSs were sufficient to confer a strong nuclear localisation (Figure 3).

The canonical KRXR NLS is strongly enriched in nucleolar and nucleoplasmic genes – found in ∼50% – however presence of KRXR alone is not a good predictor of a nuclear localisation.

### A high proportion of positively charged amino acids is associated with nucleolar localisation

Motif analysis did not identify any statistically significant linear protein motifs associated with nucleolar localisation. To investigate the properties of nucleolar proteins which might confer a general mechanism for nucleolar targeting, we analysed general protein features of nucleolar proteins in comparison to nucleoplasmic proteins, nuclear (either nucleolar or nucleoplasmic) proteins and cytoplasmic proteins.

Analysis of amino acid composition showed nucleolar proteins tended to have a similar proportion of polar amino acids, fewer hydrophobic amino acids and more charged amino acids in comparison to cytoplasmic proteins (Figure 4A), while there was no bias in molecular weight (Figure 4B). Further investigating the charged amino acids, nucleolar proteins were made up of a similar proportion of negatively charged (acidic) amino acids to cytoplasmic proteins but had far more positively charged (basic) amino acids, and correspondingly tended to have higher isoelectric points (Figure 4C).

**Figure 4.**
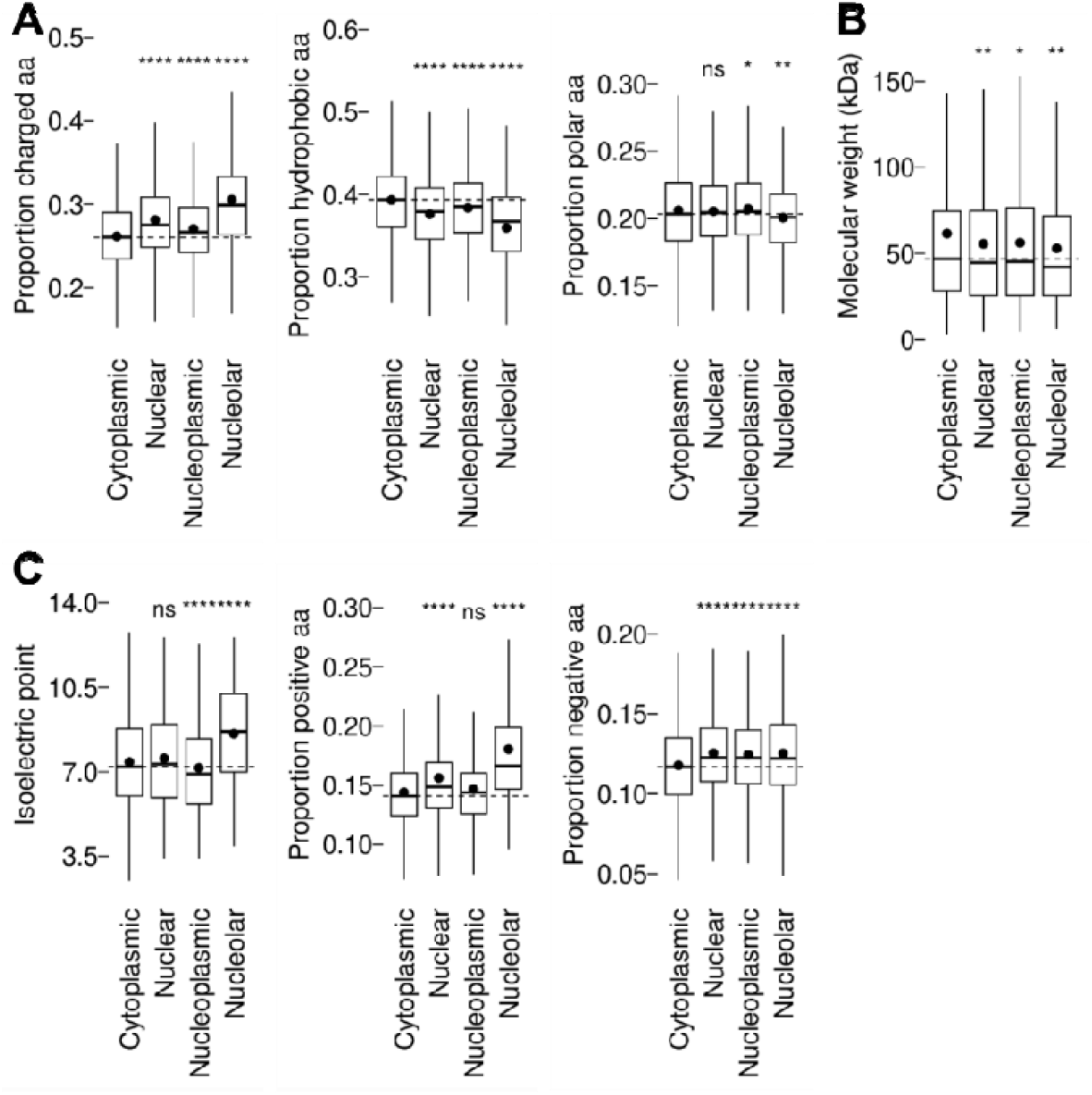
Properties of *T. brucei* nuclear, nucleoplasmic and nucleolar proteins. A. Proportion of charged (RHKDE), hydrophobic (AILMFWYV) or polar (STNQ) amino acids found in cytoplasmic, nuclear, nucleoplasmic or nucleolar proteins, as classified by the cut-offs indicated in Figure 2. B. Molecular weights of cytoplasmic, nuclear, nucleoplasmic or nucleolar proteins. C. Exploration of the abundance of charged amino acids in nucleolar genes shown in Fig. 2A. Isoelectric point and proportion of positively or negatively charged amino acids of cytoplasmic, nuclear, nucleoplasmic or nucleolar proteins. Statistical significance was assessed using the Wilcoxon signed-rank test (ns not significant, * p≤0.05, ** p≤0.01, *** p≤0.001, **** p≤0.0001).

Three other eukaryotes have comparable genome-wide protein localisation resources to *T. brucei*: the yeast species *Saccharomyces cerevisiaei* and *Schizosaccharomyces pombe* (by protein tagging) and human cell lines (by antibody). We used these to determine if the tendency for nucleolar proteins to be highly positively charged was conserved across these species using an equivalent analysis (Fig. S1). As genome-wide quantitative analysis of nucleolar/nucleoplasm partition is not available for these species we used the manually assigned annotation terms from each localisation project to determine if a protein was nucleolar. This showed that in each organism nucleolar proteins tend to have many charged amino acids and disproportionately many positively charged amino acids, although *S. cerevisiae* and *S. pombe* also had increased numbers of negatively charged amino acids in nucleolar proteins too.

### Positively charged (basic) amino acids are sufficient for nucleolar targeting

mNG is a neutral (isoelectric point 7.2) globular protein with a typical proportion of negatively (25/237, 10.5%) and positively (32/237, 13.5%) charged amino acids, which localises throughout the cytoplasm and nucleus when expressed in *T. brucei* (Figure 5A). Fusion with an NLS confers a nuclear localisation with the protein present in both the nucleoplasm and the nucleolus (Figure 3). We asked whether a shift towards more nucleolar protein-like properties (i.e. a greater proportion of charged, particularly positively charged, amino acids) could confer a nucleolar localisation.

**Figure 5.**
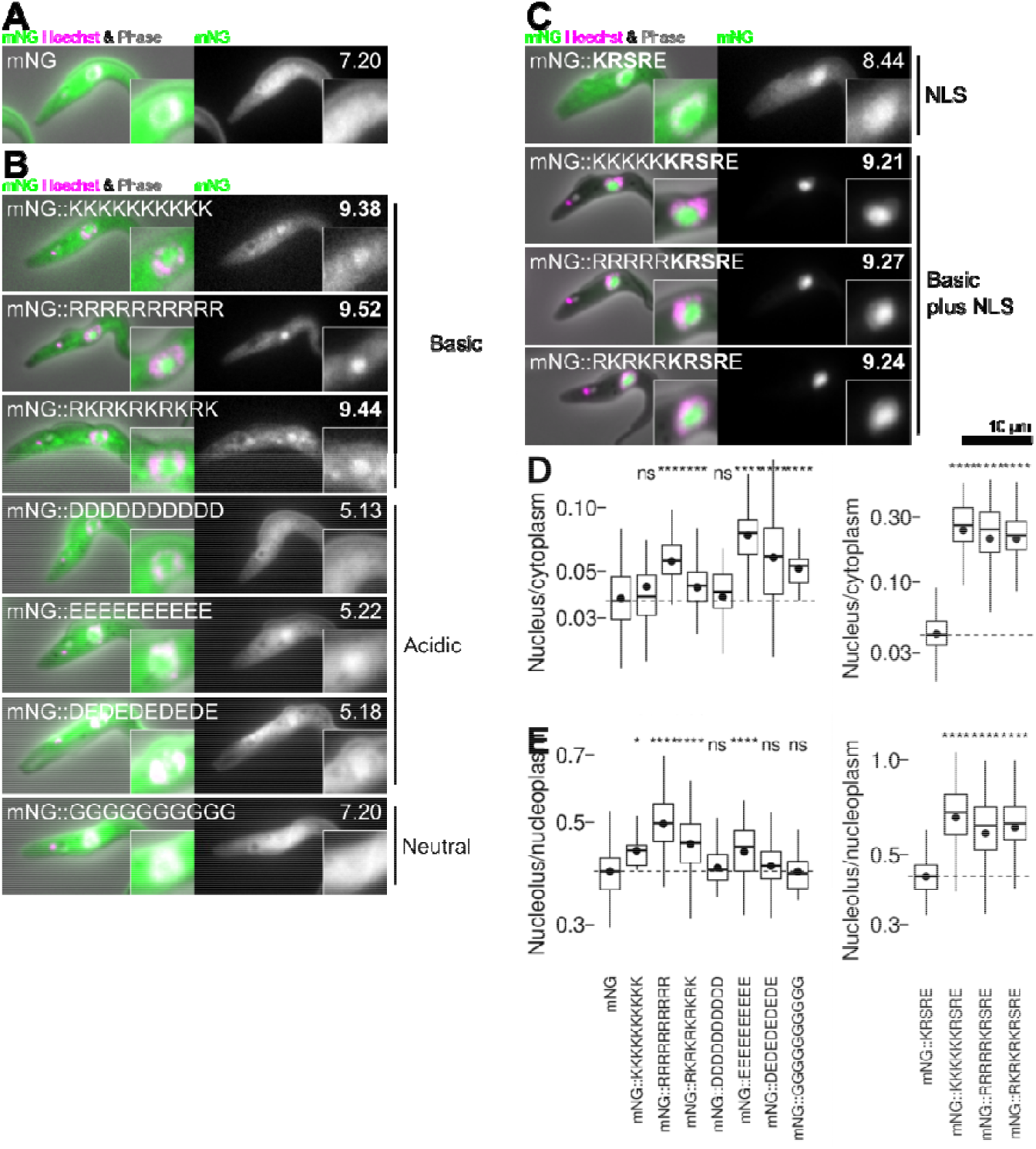
Basic sequences are sufficient for nucleolar targeting and targeting is enhanced with an NLS. A. The localisation of the fluorescent reporter protein mNG expressed from the tubulin locus. For each cell line, the mNG fusion protein pI is shown in the top right, in bold if >8.50. B. The localisation of mNG with 10 amino acid runs of basic (K or R), acidic (D or E) or neutral (G) amino acids fused to the C terminus. C. The localisation of mNG with 5 basic amino acids (K, R or a mixture) and an NLS (KRSRE) fused to the C terminus. D. Plots of automated quantitation of the nucleus/cytoplasm mNG fluorescence signal partition from the cell lines in A-C. E. As for D, but plotting the nucleolus/nucleoplasm mNG fluorescence signal partition. Statistical significance was assessed using the Wilcoxon signed-rank test (ns not significant, * p≤0.05, ** p≤0.01, *** p<0.001, **** p≤0.0001).

Fusion of mNG with short positively charged (KKKKKKKKKK, RRRRRRRRRR or RKRKRKRKR) sequences conferred a clear nucleolar localisation. Much protein remained in the cytoplasm, however of the protein in the nucleus there was a high nucleolus/nucleoplasm partition (Figure 5B). In contrast, fusion with a neutral (GGGGGGGGGG) sequence did not. Fusion with negatively charged (DDDDDDDDDD, EEEEEEEEEE, DEDEDEDEDE) gave some nuclear enrichment, particularly for E containing sequences. However clear nucleolar targeting/nucleoplasmic exclusion did not occur (Figure 5B).

A notable basic motif, RG or RGG degenerate repeats, have been associated with nucleolar localisation, notably FIB1. The *T. brucei* ortholog, NOP1, also has an RG-rich N terminal domain. However, *T. brucei* also has three NOP1 paralogs and one has all but one RG truncated from the N terminus. Fewer RGs corresponds to significantly weaker partition to the nucleolus (Fig. S5).

These terminal runs of charged amino acids are unlike many proteins; charged resides tend to be more dispersed. To test whether charge distributed within a protein can also confer nucleolar targeting we exploited the natural protein targeting systems of cells. The mitochondrion and glycosome (modified peroxisomes) have N and C terminal targeting sequences respectively – tagging by fusing mNG to the N terminus of many mitochondrial proteins prevents localisation to the mitochondrion, similarly C terminal tagging can disrupt glycosomal protein localisation (Fig. S3). We analysed all mitochondrial (Fig. S3B) and glycosomal (Fig. S3A) proteins in the TrypTag data set which gave a cytoplasmic, nuclear or nucleolar mislocalisation when their targeting sequence was disrupted by tagged at the N or C terminus respectively. Of these 26, 10 localised to the nucleolus. Of the 11 proteins with predicted pI >8.5, 8 localised to the nucleolus, a strong enrichment (p<10^−5^, Chi squared test). Mitochondrial and glycosomal proteins are normally separated with a membrane from the cytoplasm/nucleoplasm/nucleolus and should have no specific interactions with nuclear proteins, therefore a physicochemical property like pI is plausibly involved.

Finally, we further investigated short strongly negatively charged sequences, motivated by their ambiguous effect when fused to mNG (Figure 5) and tendency to arise as nucleolar-enriched motifs (although not statistically significant). We identified two proteins (Tb927.10.2310 and Tb927.9.1560) which localise to the nucleolus with a single DE rich sequence near the C terminus of the protein (Fig. S4A). Truncation to remove the acidic sequence (SADDDDDDVEIPEIDMED and SEEEEEEEEPSFEETSSDDDD respectively) did not prevent nucleolar targeting of either gene, and fusing 10 amino acids of these sequences to mNG did not confer a nucleolar localisation. In fact in both cases, truncation to remove the acidic sequence slightly, although statistically significantly, increased partition to the nucleolus (Fig. S4B).

### Functional consequence of targeting

Through its function in ribosome assembly, the largest flux of protein into the nucleolus is likely ribosome proteins into the granular compartment. However, cells face a challenge – most eukaryotes including *T. brucei* assemble at least two types of ribosomes, the mitochondrial ribosome (mitoribosome) in addition to the cytoplasmic ribosome (cytoribosome). We asked whether the mitoribosome proteins had fewer positively charged residues. In humans, yeast and *T. brucei*, cytoribosome proteins have more positively charged residues than mitoribosome proteins (Figure 6A). The same result was obtained with the first 30 amino acids trimmed from the mitoribosome protein N terminus, confirming that presence of an N terminal mitochondrial targeting signal is not responsible. A high proportion of positively charged residues was also seen in the ribosomes of eukaryotes with greatly reduced mitochondria lacking mitoribosomes, archaea ribosomes (the closest prokaryote relative of the eukaryote cytoribosome) and alphaproteobacter (the closest prokaryote relative of the eukaryote mitoribosome) (Figure 6B), consistent with the low proportion of positively charged residues in mitoribosomes proteins being an adaptation to reduce partition of the nucleolus. The distinct amino acid composition of cytoribosome and mitoribosome proteins can be visualised by t-distributed stochastic neighbour embedding (t-SNE) of the proportion of each amino acid in the sequence. A subset of nucleolar proteins has similar overall amino acid compositions (Figure 6C), and most cytoribosome proteins fall in this cluster while mitoribosome proteins cluster elsewhere (Figure 6D).

**Figure 6.**
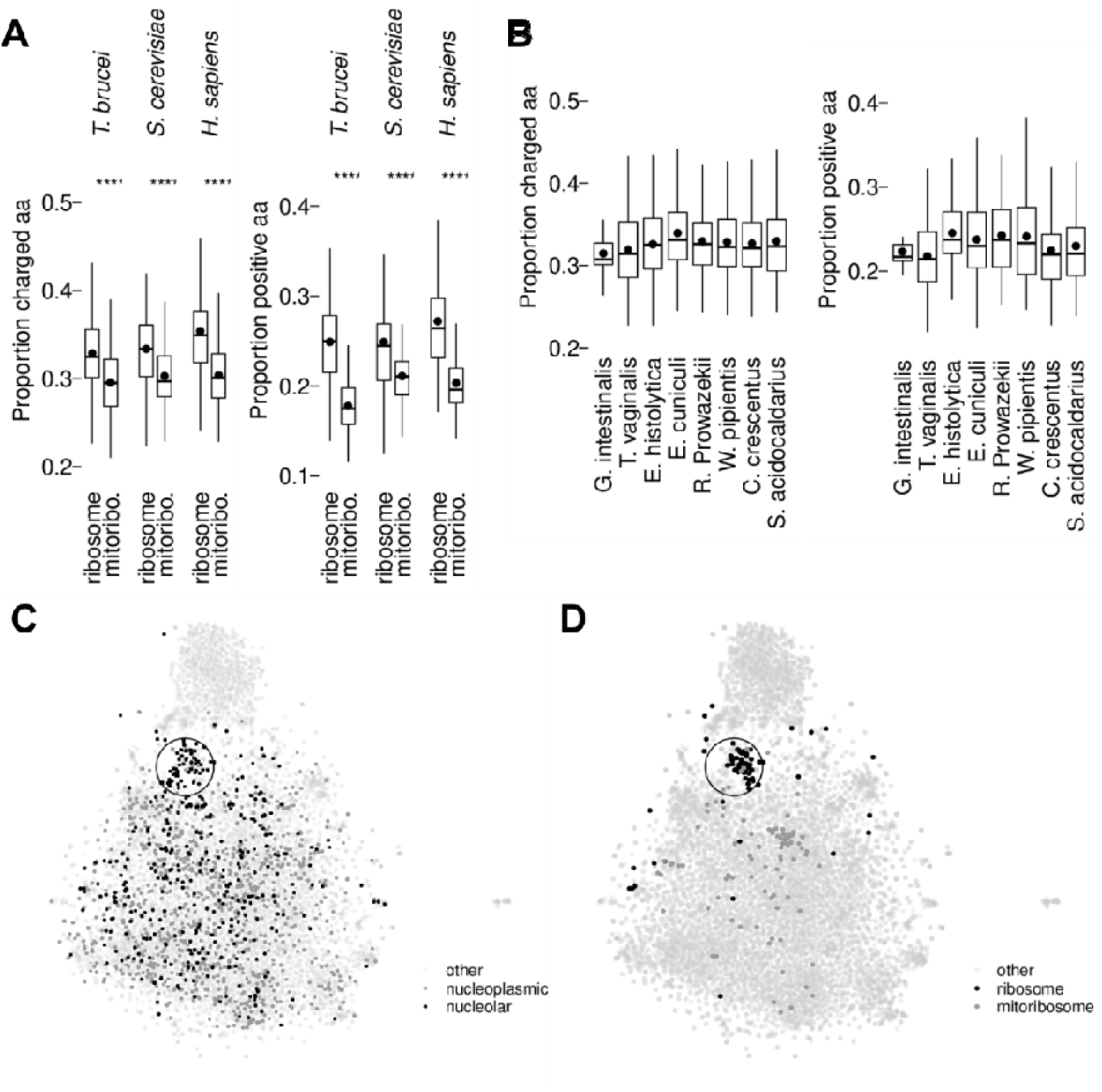
Many nucleolar proteins have similar amino acid composition to ribosome proteins but not mitochondrial ribosome proteins. A. Proportion of charged (RHKDE) or positively charged (KRH) amino acids found in *T. brucei, S. cerevisiae* and *H. sapiens* ribosome and mitochondrial ribosome (mitoribo.) proteins. B. Proportion of charged or positively charged amino acids found in ribosome proteins of organisms which do not have mitochondrial ribosomes, either because they are eukaryotes which have heavily reduced mitochondria *(G. i*., *T. v*., *E. h*., *E. c*.), they are archaea *(R. p*., *W. p*.) or they are bacteria (alphaproteobacteria, *C. c*., *S. c*.). C. t-SNE plot of all *T. brucei* proteins by amino acid composition (proportion of sequence made up of each residue), highlighting nucleoplasmic and nucleolar proteins. A subset of nucleolar proteins lies in a cluster (circled). D. The same plot as A, but highlighting ribosome and mitoribosome proteins.

## Discussion

The nucleus is an ancient organelle and, as expected, much of its molecular cell biology is conserved across all eukaryotes. As an early-branching species, *T. brucei* are informative for identifying these ancestral features. Of the nuclear import machinery, many components are conserved (albeit with significant adaptations)(Canela-Pérez et al., 2019; Keminer and Peters, 1999; Mattaj and Englmeier, 1998; Obado et al., 2016), as is the monopartite KRXR NLS(Marchetti et al., 2000). Our analysis, using genome-wide protein localisation data to quantify nucleolar enrichment, indicated basicity from both short basic IDRs or basic residues throughout the protein, and therefore protein charge, is the conserved feature key for nucleolar targeting.

Our *de novo* search for linear motifs readily re-revealed the KRXR NLS (Figure 2, Figure 3) but did not reveal a NoLS motif. It instead pointed to the importance of the number of basic residues. Our equivalent analysis of existing genome-wide protein localisation data in humans and yeast (Fig. S1) also showed this pattern. Short linear (poly-R and poly-K) and mixed (poly-RK) sequences were sufficient for nucleolar targeting in *T. brucei* (Figure 5). This is very similar to previous analysis in mammalian cells(Martin et al., 2015; Musinova et al., 2015). We also showed that proteins with dispersed positive charge tend to mislocalise to the nucleolus when their normal targeting sequences are disrupted (Fig. S3). This indicates net charge rather than a linear motif is sufficient for nucleolar targeting, and points to basicity being the conserved feature across divergent eukaryotes for nucleolar targeting. However, positive charge alone is a very general feature and is poorly predictive. The nucleolus is also a complex structure, and this does not address targeting to known nucleolar subcompartments.

While charge alone appeared sufficient for some nucleolar targeting, *T. brucei* nucleolar proteins, as in humans(Stenström et al., 2020), tended to have larger IDRs. The basic charge and low hydrophobicity typical among *T. brucei* nucleolar proteins are associated with condensate formation and IDRs(Quiroz and Chilkoti, 2015; Uversky, 2002). The Das-Pappu diagram of states for polyampholytic IDRs(Das et al., 2015; Holehouse et al., 2017) showed *T. brucei* nucleolar proteins are often predicted to be polyampholytic coils and hairpins (Fig. S2). Comparable polymer physics systems indicate this can be favourable for phase separation(Bianchi et al., 2020; Srivastava and Muthukumar, 1996), speaking to the wider question of how the *T. brucei* nucleolus forms.

Nucleolar assembly carries additional importance in *T. brucei* as, in mammalian infective life cycle stages, they form a second distinct RNA Pol I nuclear compartment called the expression site body (ESB)(Navarro and Gull, 2001b). The ESB is vital for antigenic variation. It has no function in rRNA transcription or ribosome assembly but does share some components with the nucleolus (RNA Pol I and basal transcription factors)(Nguyen et al., 2012; Nguyen et al., 2014) and has some unique components(Escobar et al., 2021; Faria et al., 2019). Its concurrent existence with the nucleolus means a distinction to their sorting mechanisms exists. How proteins are sorted to the ESB vs nucleolus is therefore an important question for the future.

Protein charge being responsible for nucleolar partition is consistent with LLPS models for nucleolus formation. Some metazoan nucleolar proteins, notably FIB1 and NPM1, can phase separate in vitro and form mutually immiscible condensates which mimic nucleolar compartments(Bianchi et al., 2020; Haynes et al., 2006). NPM1 is a major component of the granular compartment in metazoa with a series of negatively-charged acidic tracts in its IDRs. Proteins with characterised basic tracts which act as NoLSs, including APE1 and ARF, have recently been proposed to partition to the nucleolus through the interaction of their basic R motifs with the acidic tracts of NPM1(Lindström and Zhang, 2006; Lirussi et al., 2012; López et al., 2020; Mitrea et al., 2016; Mitrea et al., 2018). Based on our evidence in *T. brucei*, we argue charge interactions with the granular component phase is the general nucleolar targeting phenomenon across eukaryotes.

*T. brucei* nucleolar architecture is incompletely described, but, like most eukaryotes, includes granular (ribosome assembly) and Pol I (transcription) compartments(Daniels et al., 2010). *T. brucei* does not have a clear ortholog of NPM1, although multiple nucleolar proteins with similar acidic tracts and a high proportion of charged amino acids are present. It does, however, have multiple orthologs of FIB1 (called NOP1), which also contain multiple RG motifs that contribute to partition to the nucleolus (Fig. S5). This is strongly associated with LLPS, as shown by LAF-1 (*C. elegans* P granules) and DDX4, which have RG motifs necessary for phase separation(Elbaum-Garfinkle et al., 2015; Nott et al., 2015). However, tentatively, NOP1 localises to smaller nucleolar subdomains and we suspect our analysis is dominated by partition to the larger granular compartment, therefore relating most strongly to ribosome assembly.

We identified a peculiar feature of mitoribosome proteins, that they have a lower proportion of basic amino acids than cytoplasmic ribosome proteins, despite basic amino acids often being common in nucleic acid-interacting proteins and proteins in ribonuclear complexes. This is not a peculiarity of *T. brucei* and is also the case in humans and yeast (Figure 6). To the best of our knowledge this has not previously been noted and suggests a selection pressure for mitoribosome proteins to be less basic. Evolution of the mitoribosome is complex, having undergone extensive remodelling during its evolutionary course(Ku et al., 2015) after acquisition of the mitochondrion by endosymbiosis of an α-proteobacteria by an ancestral eukaryote(Gray, 2017). This includes acquiring N-terminal mitochondrial localisation signals known as presequences. These presequences are known to be positively charged(Dudek et al., 2013), and despite this the overall number of mitoribosome protein positively charged residues is still lower than cytoribosome proteins, further implicating a selection pressure. Given that mitochondrial proteins do not generally enter the mitochondrion co-translationally, they necessarily spend some time in the cytoplasm and would have the opportunity to enter the nucleus by accident. The less basic nature of mitoribosome proteins would, therefore, help prevent their partition to the nucleolus. However, we cannot exclude a selection pressure to assist transport across the double mitochondrial membranes.

We saw that many basic mitochondrial proteins can mislocalise to the nucleolus when the mitochondrial localisation signal is disrupted by N terminal tagging (Fig. S3). While this indicates that mitochondrial targeting is sufficient to overcome nucleolar targeting arising from protein physicochemical properties, mitoribosome protein targeting would certainly be aided by a mechanism that redirects proteins away from the nucleolus. It may also prevent interference of mitoribosome proteins with cytoribosome assembly in the nucleolus.

## Conclusion

Proteins with a large number of basic residues, low hydrophobicity and high intrinsic disorder are common in *T. brucei* nucleolar proteins, with basic tracts or an overall basic nature sufficient for nucleolar targeting. Together, this is consistent with LLPS models for nucleolar formation and partition of proteins to the compartment. As *T. brucei* is an early-branching eukaryote, and similar features have been implicated in nucleolar targeting in other organisms, this mechanism of nucleolar targeting is likely conserved across eukaryotes. As mitoribosome proteins have a more acidic sequence than cytoribosome proteins, this contributes to cytoribosome vs mitoribosome protein sorting.

## Methods

### Cells and cell culture

Procyclic form *Trypanosoma brucei brucei* strain TREU927 were used as they were used for the original *T. brucei*(Berriman et al., 2005) genome and the TrypTag genome-wide protein localisation project(Dean et al., 2017). They were grown in SDM-79(Brun and Schönenberger, 1979) at 28°C, and maintained between approximately 6×10^5^ and 2×10^7^ cells/ml by regular subculture.

### Genetic modification

Cell lines stably expressing proteins tagged at the N or C terminus with the fluorescent protein mNeonGreen (mNG)(Shaner et al., 2013) were generated by modification of one of the endogenous alleles. Tagging was carried out as previously described, using long primer PCR using the plasmid pPOT v4 BLAST mNG as the template to generate tagging constructs. The template plasmid provides a standard fluorescent protein and drug selection marker coding sequences, while forward and reverse long primers introduce gene-specific 80 bp 5’ and 3’ homology arms – to either the 5’ UTR and start of the target gene ORF or the end of the target gene ORF (excluding the stop codon) and the 3’ UTR, for N and C terminal tagging respectively(Dean et al., 2015). High throughput electroporation was used to transfect *T. brucei* with the tagging constructs, which integrate into the target locus by homologous recombination(Dyer et al., 2016). 10 µg/ml (PCF) Blasticidin S Hydrochloride used to select for successful transfectants.

Cell lines stably expressing truncated tagged proteins were generated as for tagging except with shifted homology within the target gene ORF to introduce a truncation at the tagged terminus, as previously described(Dean et al., 2015). For N terminal truncation, homology to the target gene ORF in the reverse primer was shifted the necessary number of codons into the start of the ORF and for C terminal truncation homology to the target gene ORF in the forward primer was shifted the necessary number of codons into the end of the ORF.

Cell lines expressing mNG with a N or C terminal candidate targeting sequence were also generated using similar a PCR-based method. Here, the homology arms were designed such that one copy of α-tubulin in the multi-copy tubulin array is replaced by the mNG and drug selection marker coding sequences. Using the standard pPOT primer binding sites(Dean et al., 2015), a candidate targeting sequence of up to 10 codons can be fused to the mNG coding sequence, using 50 bp homology and 30 bp encoding the targeting sequence on the forward or reverse primer for introduction to the C or N terminus of mNG respectively.

### Microscopy

Live cells were stained with Hoechst 33342 and adhered, live, to glass slides as previously described(Dean and Sunter, 2020). mNG and Hoechst 33342 fluorescence and phase contrast micrographs were captured on the same microscope and using identical settings as the TrypTag genome wide protein localisation project, a DM5500 B (Leica) upright widefield epifluorescence microscope using a plan apo NA/1.4 63× phase contrast oil immersion objective (Leica, 15506351) and a Neo v5.5 (Andor) sCMOS camera using MicroManager(Edelstein et al., 2010).

### Automated image analysis

Image analysis builds on our previous approaches(Wheeler, 2020; Wheeler et al., 2012) using ImageJ(Collins, 2007). All images analysed were at 0.103 μm/px. They were first flat-field corrected by subtracting the median of all images captured on a particular day. To identify cells, phase contrast images were pre-processed by sequential Gaussian unsharp filters with radii from 1 to 35 px at 5 px steps with 0.4 weight, then an intensity threshold of the image mean minus 1× standard deviation was applied. Cells were taken as objects between 2000 and 7000 px^2^, with a minimum pixel value at least two standard deviations under the mean.

To identify nuclei, Hoechst 33342 fluorescence images were pre-processed by a 1 px radius Gaussian blur and a rolling ball subtraction with radius 15 px. Local maxima with prominence over 1.5× image standard deviation were taken as DNA containing objects, with a threshold equal to 0.4× the local maxima. *T. brucei* have two DNA-containing structures, the nucleus (N) and kinetoplast (K) and K divides before N in the cell cycle, therefore in cells with two DNA-containing structures (expected to be 1K1N) the largest was the taken as the nucleus, in three structure cells (expected to be 2K1N) the largest was taken as a nucleus and four structure cells (expected to be 2K2N) the larges two were taken as nuclei. Mean nucleus radius *r* was taken as the average of the major and minor axes of an ellipse fitted to the thresholded object. Nucleoli appear as small circular regions of lower Hoechst 33342 in the nucleus. To identify nucleoli, the darkest point within the nucleus at least *r*/4 from its edges was taken as the nucleolar centre and assumed to have a radius *r*/4.

Nuclear/cytoplasm partition was taken as the ratio of mean nuclear signal to mean cytoplasmic signal in the mNG fluorescence channel, nucleolar/nucleus partition was taken as the ratio of mean nucleolar signal to mean nucleoplasm (ie. excluding the nucleolus) signal. Data for the TrypTag genome tagging project dataset represent mean partition for all cells (typically >200), plotting N and C terminally tagged cell lines separately. Other data was further filtered to exclude cells not expressing the fluorescently tagged protein, sample sizes indicate the number of cells.

### Protein primary sequence analysis

Meme(Bailey et al., 2009) version 5.1.1 was used to identify linear motifs enriched in nuclear and nucleolar proteins, searching for motifs with one occurrence per sequence and widths between 4 and 16. IDRs were identified using IUPred2A(Erdős and Dosztányi, 2020; Mészáros et al., 2018), taking residues with a score over 0.5 as disordered.

Human protein localisations were taken from the Human Cell Atlas (accessed Dec 2020), taking any proteins annotated with terms including “nucleoli” as nucleolar, and proteins annotated with any nuclear lumen structures as a nuclear(Thul et al., 2017). Yeast localisations were taken from the Yeast GFP Fusion Localization Database (accessed Dec 2020), using their nucleolar and nuclear annotations(Huh et al., 2003).

*T. brucei* protein sequences were taken from TriTrypDB v51(Aslett et al., 2010), for all other species protein sequences were taken from UniProt. *T. brucei* cytoribosome and mitoribosome protein lists were derived from those identified by affinity purification and/or cryoelectron microscopy structures(Hashem et al., 2013; Saurer et al., 2019; Zíková et al., 2008). In other species, lists were derived from UniProt protein annotations – for example “60S ribosomal protein LX” or “40S ribosomal protein SX” for human cytoribosomes.

## Abbreviations

LLPS: liquid-liquid phase separation
NLS: nuclear localisation signal
NoLS: nucleolar localisation signal
Pol I: RNA Polymerase I
IDR: intrinsically disordered region
mNG: mNeonGreen

## Notes

### Competing Interest Statement

The authors have declared no competing interest.

## References

Afrin, M., Kishmiri, H., Sandhu, R., Rabbani, M. a. G. and Li, B. (2020). Trypanosoma brucei RAP1 Has Essential Functional Domains That Are Required for Different Protein Interactions. mSphere 5,.

Aslett, M., Aurrecoechea, C., Berriman, M., Brestelli, J., Brunk, B. P., Carrington, M., Depledge, D. P., Fischer, S., Gajria, B., Gao, X., et al. (2010). TriTrypDB: a functional genomic resource for the Trypanosomatidae. Nucleic Acids Res 38, D457–D462.

Bailey, T. L., Boden, M., Buske, F. A., Frith, M., Grant, C. E., Clementi, L., Ren, J., Li, W. W. and Noble, W. S. (2009). MEME SUITE: tools for motif discovery and searching. Nucleic Acids Res 37, W202–208.

Banani, S. F., Lee, H. O., Hyman, A. A. and Rosen, M. K. (2017). Biomolecular condensates: organizers of cellular biochemistry. Nat. Rev. Mol. Cell Biol. 18, 285–298.

Berriman, M., Ghedin, E., Hertz-Fowler, C., Blandin, G., Renauld, H., Bartholomeu, D. C., Lennard, N. J., Caler, E., Hamlin, N. E., Haas, B., et al. (2005). The genome of the African trypanosome Trypanosoma brucei. Science 309, 416–22.

Bianchi, G., Longhi, S., Grandori, R. and Brocca, S. (2020). Relevance of Electrostatic Charges in Compactness, Aggregation, and Phase Separation of Intrinsically Disordered Proteins. Int J Mol Sci 21,.

Boeynaems, S., Alberti, S., Fawzi, N. L., Mittag, T., Polymenidou, M., Rousseau, F., Schymkowitz, J., Shorter, J., Wolozin, B., Van Den Bosch, L., et al. (2018). Protein Phase Separation: A New Phase in Cell Biology. Trends Cell Biol 28, 420–435.

Brangwynne, C. P., Eckmann, C. R., Courson, D. S., Rybarska, A., Hoege, C., Gharakhani, J., Jülicher, F. and Hyman, A. A. (2009). Germline P granules are liquid droplets that localize by controlled dissolution/condensation. Science 324, 1729–1732.

Brangwynne, C. P., Mitchison, T. J. and Hyman, A. A. (2011). Active liquid-like behavior of nucleoli determines their size and shape in Xenopus laevis oocytes. Proc. Natl. Acad. Sci. U.S.A. 108, 4334–4339.

Brun, R. and Schönenberger, M. (1979). Cultivation and in vitro cloning or procyclic culture forms of Trypanosoma brucei in a semi-defined medium. Short communication. Acta Trop 36, 289–292.

Canela-Pérez, I., López-Villaseñor, I., Cevallos, A. M. and Hernández, R. (2018). Nuclear distribution of the Trypanosoma cruzi RNA Pol I subunit RPA31 during growth and metacyclogenesis, and characterization of its nuclear localization signal. Parasitol Res 117, 911–918.

Canela-Pérez, I., López-Villaseñor, I., Mendoza, L., Cevallos, A. M. and Hernández, R. (2019). Nuclear localization signals in trypanosomal proteins. Mol Biochem Parasitol 229, 15–23.

Canela-Pérez, I., López-Villaseñor, I., Cevallos, A. M. and Hernández, R. (2020). Trypanosoma cruzi Importin α: ability to bind to a functional classical nuclear localization signal of the bipartite type. Parasitol Res 119, 3899–3907.

Chelsky, D., Ralph, R. and Jonak, G. (1989). Sequence requirements for synthetic peptide-mediated translocation to the nucleus. Mol Cell Biol 9, 2487–2492.

Collins, T. J. (2007). ImageJ for microscopy. BioTechniques 43, 25–30.

Daniels, J.-P., Gull, K. and Wickstead, B. (2010). Cell biology of the trypanosome genome. Microbiol. Mol. Biol. Rev. 74, 552–569.

Das, R. K., Ruff, K. M. and Pappu, R. V. (2015). Relating sequence encoded information to form and function of intrinsically disordered proteins. Curr. Opin. Struct. Biol. 32, 102–112.

Dean, S. and Sunter, J. (2020). Light Microscopy in Trypanosomes: Use of Fluorescent Proteins and Tags. Methods Mol. Biol. 2116, 367–383.

Dean, S., Sunter, J., Wheeler, R. J., Hodkinson, I., Gluenz, E. and Gull, K. (2015). A toolkit enabling efficient, scalable and reproducible gene tagging in trypanosomatids. Open Biol 5, 140197.

Dean, S., Sunter, J. D. and Wheeler, R. J. (2017). TrypTag.org: A Trypanosome Genome-wide Protein Localisation Resource. Trends in Parasitology 33, 80–82.

Dingwall, C., Sharnick, S. V. and Laskey, R. A. (1982). A polypeptide domain that specifies migration of nucleoplasmin into the nucleus. Cell 30, 449–458.

Duan, T.-L., He, G.-J., Hu, L.-D. and Yan, Y.-B. (2019). The Intrinsically Disordered C-Terminal Domain Triggers Nucleolar Localization and Function Switch of PARN in Response to DNA Damage. Cells 8,.

Dudek, J., Rehling, P. and van der Laan, M. (2013). Mitochondrial protein import: common principles and physiological networks. Biochim Biophys Acta 1833, 274–285.

Dyer, P., Dean, S. and Sunter, J. (2016). High-throughput Gene Tagging in Trypanosoma brucei. Journal of Visualized Experiments e54342.

Edelstein, A., Amodaj, N., Hoover, K., Vale, R. and Stuurman, N. (2010). Computer control of microscopes using µManager. Curr Protoc Mol Biol Chapter 14, Unit14.20.

Elbaum-Garfinkle, S., Kim, Y., Szczepaniak, K., Chen, C. C.-H., Eckmann, C. R., Myong, S. and Brangwynne, C. P. (2015). The disordered P granule protein LAF-1 drives phase separation into droplets with tunable viscosity and dynamics. Proc Natl Acad Sci U S A 112, 7189–7194.

Erdős, G. and Dosztányi, Z. (2020). Analyzing Protein Disorder with IUPred2A. Current Protocols in Bioinformatics 70, e99.

Escobar, L. L., Hänisch, B., Halliday, C., Dean, S., Sunter, J. D., Wheeler, R. J. and Gull, K. (2021). Monoallelic antigen expression in trypanosomes requires a stage-specific transcription activator.

Faria, J., Glover, L., Hutchinson, S., Boehm, C., Field, M. C. and Horn, D. (2019). Monoallelic expression and epigenetic inheritance sustained by a Trypanosoma brucei variant surface glycoprotein exclusion complex. Nat Commun 10, 3023.

Feng, Z., Chen, X., Wu, X. and Zhang, M. (2019). Formation of biological condensates via phase separation: Characteristics, analytical methods, and physiological implications. J Biol Chem 294, 14823–14835.

Feric, M., Vaidya, N., Harmon, T. S., Mitrea, D. M., Zhu, L., Richardson, T. M., Kriwacki, R. W., Pappu, R. V. and Brangwynne, C. P. (2016). Coexisting Liquid Phases Underlie Nucleolar Subcompartments. Cell 165, 1686–1697.

Gomes, E. and Shorter, J. (2019). The molecular language of membraneless organelles. J. Biol. Chem. 294, 7115–7127.

Goos, C., Dejung, M., Janzen, C. J., Butter, F. and Kramer, S. (2017). The nuclear proteome of Trypanosoma brucei. PLoS ONE 12, e0181884.

Gray, M. W. (2017). Lynn Margulis and the endosymbiont hypothesis: 50 years later. Mol Biol Cell 28, 1285–1287.

Günzl, A., Bruderer, T., Laufer, G., Schimanski, B., Tu, L.-C., Chung, H.-M., Lee, P.-T. and Lee, M. G.-S. (2003). RNA polymerase I transcribes procyclin genes and variant surface glycoprotein gene expression sites in Trypanosoma brucei. Eukaryot Cell 2, 542–551.

Hashem, Y., des Georges, A., Fu, J., Buss, S. N., Jossinet, F., Jobe, A., Zhang, Q., Liao, H. Y., Grassucci, R. A., Bajaj, C., et al. (2013). High-resolution cryo-electron microscopy structure of the Trypanosoma brucei ribosome. Nature 494, 385–389.

Haynes, C., Oldfield, C. J., Ji, F., Klitgord, N., Cusick, M. E., Radivojac, P., Uversky, V. N., Vidal, M. and Iakoucheva, L. M. (2006). Intrinsic disorder is a common feature of hub proteins from four eukaryotic interactomes. PLoS Comput Biol 2, e100.

Holehouse, A. S., Das, R. K., Ahad, J. N., Richardson, M. O. G. and Pappu, R. V. (2017). CIDER: Resources to Analyze Sequence-Ensemble Relationships of Intrinsically Disordered Proteins. Biophysical Journal 112, 16–21.

Huh, W.-K., Falvo, J. V., Gerke, L. C., Carroll, A. S., Howson, R. W., Weissman, J. S. and O’Shea, E. K. (2003). Global analysis of protein localization in budding yeast. Nature 425, 686–691.

Iyama, T., Okur, M. N., Golato, T., McNeill, D. R., Lu, H., Hamilton, R., Raja, A., Bohr, V. A. and Wilson, D. M. (2018). Regulation of the Intranuclear Distribution of the Cockayne Syndrome Proteins. Sci Rep 8, 17490.

Keminer, O. and Peters, R. (1999). Permeability of single nuclear pores. Biophys J 77, 217–228.

Ku, C., Nelson-Sathi, S., Roettger, M., Sousa, F. L., Lockhart, P. J., Bryant, D., Hazkani-Covo, E., McInerney, J. O., Landan, G. and Martin, W. F. (2015). Endosymbiotic origin and differential loss of eukaryotic genes. Nature 524, 427–432.

Lafontaine, D. L. J., Riback, J. A., Bascetin, R. and Brangwynne, C. P. (2021). The nucleolus as a multiphase liquid condensate. Nat Rev Mol Cell Biol 22, 165–182.

Landeira, D. and Navarro, M. (2007). Nuclear repositioning of the VSG promoter during developmental silencing in Trypanosoma brucei. J. Cell Biol. 176, 133–139.

Li, P., Banjade, S., Cheng, H.-C., Kim, S., Chen, B., Guo, L., Llaguno, M., Hollingsworth, J. V., King, D. S., Banani, S. F., et al. (2012). Phase transitions in the assembly of multivalent signalling proteins. Nature 483, 336–340.

Lin, Y., Protter, D. S. W., Rosen, M. K. and Parker, R. (2015). Formation and Maturation of Phase-Separated Liquid Droplets by RNA-Binding Proteins. Mol. Cell 60, 208–219.

Lin, Y.-H., Forman-Kay, J. D. and Chan, H. S. (2018). Theories for Sequence-Dependent Phase Behaviors of Biomolecular Condensates. Biochemistry 57, 2499–2508.

Lindström, M. S. and Zhang, Y. (2006). B23 and ARF: friends or foes? Cell Biochem Biophys 46, 79–90.

Lirussi, L., Antoniali, G., Vascotto, C., D’Ambrosio, C., Poletto, M., Romanello, M., Marasco, D., Leone, M., Quadrifoglio, F., Bhakat, K. K., et al. (2012). Nucleolar accumulation of APE1 depends on charged lysine residues that undergo acetylation upon genotoxic stress and modulate its BER activity in cells. Mol Biol Cell 23, 4079–4096.

López, D. J., de Blas, A., Hurtado, M., García-Alija, M., Mentxaka, J., de la Arada, I., Urbaneja, M. A., Alonso-Mariño, M. and Bañuelos, S. (2020). Nucleophosmin interaction with APE1: Insights into DNA repair regulation. DNA Repair (Amst) 88, 102809.

Marchetti, M. A., Tschudi, C., Kwon, H., Wolin, S. L. and Ullu, E. (2000). Import of proteins into the trypanosome nucleus and their distribution at karyokinesis. J. Cell. Sci. 113 (Pt 5), 899–906.

Martin, E. W. and Holehouse, A. S. (2020). Intrinsically disordered protein regions and phase separation: sequence determinants of assembly or lack thereof. Emerg Top Life Sci 4, 307–329.

Martin, E. W. and Mittag, T. (2018). Relationship of Sequence and Phase Separation in Protein Low-Complexity Regions. Biochemistry 57, 2478–2487.

Martin, R. M., Ter-Avetisyan, G., Herce, H. D., Ludwig, A. K., Lättig-Tünnemann, G. and Cardoso, M. C. (2015). Principles of protein targeting to the nucleolus. Nucleus 6, 314–325.

Mattaj, I. W. and Englmeier, L. (1998). Nucleocytoplasmic transport: the soluble phase. Annu Rev Biochem 67, 265–306.

Meng, F., Na, I., Kurgan, L. and Uversky, V. N. (2015). Compartmentalization and Functionality of Nuclear Disorder: Intrinsic Disorder and Protein-Protein Interactions in Intra-Nuclear Compartments. Int J Mol Sci 17,.

Mészáros, B., Erdős, G. and Dosztányi, Z. (2018). IUPred2A: context-dependent prediction of protein disorder as a function of redox state and protein binding. Nucleic Acids Research 46, W329–W337.

Mitrea, D. M., Cika, J. A., Guy, C. S., Ban, D., Banerjee, P. R., Stanley, C. B., Nourse, A., Deniz, A. A. and Kriwacki, R. W. (2016). Nucleophosmin integrates within the nucleolus via multi-modal interactions with proteins displaying R-rich linear motifs and rRNA. Elife 5,.

Mitrea, D. M., Cika, J. A., Stanley, C. B., Nourse, A., Onuchic, P. L., Banerjee, P. R., Phillips, A. H., Park, C.-G., Deniz, A. A. and Kriwacki, R. W. (2018). Self-interaction of NPM1 modulates multiple mechanisms of liquid-liquid phase separation. Nat Commun 9, 842.

Musinova, Y. R., Lisitsyna, O. M., Golyshev, S. A., Tuzhikov, A. I., Polyakov, V. Y. and Sheval, E. V. (2011). Nucleolar localization/retention signal is responsible for transient accumulation of histone H2B in the nucleolus through electrostatic interactions. Biochim Biophys Acta 1813, 27–38.

Musinova, Y. R., Kananykhina, E. Y., Potashnikova, D. M., Lisitsyna, O. M. and Sheval, E. V. (2015). A charge-dependent mechanism is responsible for the dynamic accumulation of proteins inside nucleoli. Biochim Biophys Acta 1853, 101–110.

Navarro, M. and Gull, K. (2001a). A pol I transcriptional body associated with VSG mono-allelic expression in Trypanosoma brucei. Nature 414, 759–763.

Navarro, M. and Gull, K. (2001b). A pol I transcriptional body associated with VSG mono-allelic expression in Trypanosoma brucei. Nature 414, 759–763.

Nguyen, T. N., Nguyen, B. N., Lee, J. H., Panigrahi, A. K. and Günzl, A. (2012). Characterization of a Novel Class I Transcription Factor A (CITFA) Subunit That Is Indispensable for Transcription by the Multifunctional RNA Polymerase I of Trypanosoma brucei. Eukaryot Cell 11, 1573–1581.

Nguyen, T. N., Müller, L. S. M., Park, S. H., Siegel, T. N. and Günzl, A. (2014). Promoter occupancy of the basal class I transcription factor A differs strongly between active and silent VSG expression sites in Trypanosoma brucei. Nucleic Acids Res. 42, 3164–3176.

Nott, T. J., Petsalaki, E., Farber, P., Jervis, D., Fussner, E., Plochowietz, A., Craggs, T. D., Bazett-Jones, D. P., Pawson, T., Forman-Kay, J. D., et al. (2015). Phase Transition of a Disordered Nuage Protein Generates Environmentally Responsive Membraneless Organelles. Mol Cell 57, 936–947.

Obado, S. O., Brillantes, M., Uryu, K., Zhang, W., Ketaren, N. E., Chait, B. T., Field, M. C. and Rout, M. P. (2016). Interactome Mapping Reveals the Evolutionary History of the Nuclear Pore Complex. PLOS Biology 14, e1002365.

Pays, E., Tebabi, P., Pays, A., Coquelet, H., Revelard, P., Salmon, D. and Steinert, M. (1989). The genes and transcripts of an antigen gene expression site from T. brucei. Cell 57, 835–845.

Quiroz, F. G. and Chilkoti, A. (2015). Sequence heuristics to encode phase behaviour in intrinsically disordered protein polymers. Nat Mater 14, 1164–1171.

Saurer, M., Ramrath, D. J. F., Niemann, M., Calderaro, S., Prange, C., Mattei, S., Scaiola, A., Leitner, A., Bieri, P., Horn, E. K., et al. (2019). Mitoribosomal small subunit biogenesis in trypanosomes involves an extensive assembly machinery. Science 365, 1144–1149.

Savada, R. P. and Bonham-Smith, P. C. (2013). Charge versus sequence for nuclear/nucleolar localization of plant ribosomal proteins. Plant Mol Biol 81, 477–493.

Sawyer, I. A., Sturgill, D. and Dundr, M. (2019a). Membraneless nuclear organelles and the search for phases within phases. Wiley Interdiscip Rev RNA 10, e1514.

Sawyer, I. A., Bartek, J. and Dundr, M. (2019b). Phase separated microenvironments inside the cell nucleus are linked to disease and regulate epigenetic state, transcription and RNA processing. Semin Cell Dev Biol 90, 94–103.

Schmidt-Zachmann, M. S. and Nigg, E. A. (1993). Protein localization to the nucleolus: a search for targeting domains in nucleolin. J Cell Sci 105 (Pt 3), 799–806.

Scott, M. S., Boisvert, F.-M., McDowall, M. D., Lamond, A. I. and Barton, G. J. (2010). Characterization and prediction of protein nucleolar localization sequences. Nucleic Acids Res. 38, 7388–7399.

Scott, M. S., Troshin, P. V. and Barton, G. J. (2011). NoD: a Nucleolar localization sequence detector for eukaryotic and viral proteins. BMC Bioinformatics 12, 317.

Shaner, N. C., Lambert, G. G., Chammas, A., Ni, Y., Cranfill, P. J., Baird, M. A., Sell, B. R., Allen, J. R., Day, R. N., Israelsson, M., et al. (2013). A bright monomeric green fluorescent protein derived from Branchiostoma lanceolatum. Nat Meth 10, 407–409.

Srivastava, D. and Muthukumar, M. (1996). Sequence Dependence of Conformations of Polyampholytes. Macromolecules 29, 2324–2326.

Stenström, L., Mahdessian, D., Gnann, C., Cesnik, A. J., Ouyang, W., Leonetti, M. D., Uhlén, M., Cuylen-Haering, S., Thul, P. J. and Lundberg, E. (2020). Mapping the nucleolar proteome reveals a spatiotemporal organization related to intrinsic protein disorder. Mol Syst Biol 16, e9469.

Thul, P. J., Åkesson, L., Wiking, M., Mahdessian, D., Geladaki, A., Blal, H. A., Alm, T., Asplund, A., Björk, L., Breckels, L. M., et al. (2017). A subcellular map of the human proteome. Science 356, eaal3321.

Uversky, V. N. (2002). Natively unfolded proteins: a point where biology waits for physics. Protein Sci 11, 739–756.

Wang, Z. and Zhang, H. (2019). Phase Separation, Transition, and Autophagic Degradation of Proteins in Development and Pathogenesis. Trends Cell Biol 29, 417–427.

Weber, S. C. (2017). Sequence-encoded material properties dictate the structure and function of nuclear bodies. Curr Opin Cell Biol 46, 62–71.

Wheeler, R. J. (2020). ImageJ for Partially and Fully Automated Analysis of Trypanosome Micrographs. In Trypanosomatids: Methods and Protocols (ed. Michels, P. A. M.), Ginger, M. L.), and Zilberstein, D.), pp. 385–408. New York, NY: Springer US.

Wheeler, R. J., Gull, K. and Gluenz, E. (2012). Detailed interrogation of trypanosome cell biology via differential organelle staining and automated image analysis. BMC Biology 10, 1.

Woodruff, J. B., Hyman, A. A. and Boke, E. (2018). Organization and Function of Non-dynamic Biomolecular Condensates. Trends Biochem Sci 43, 81–94.

Wright, P. E. and Dyson, H. J. (2015). Intrinsically disordered proteins in cellular signalling and regulation. Nat Rev Mol Cell Biol 16, 18–29.

Zíková, A., Panigrahi, A. K., Dalley, R. A., Acestor, N., Anupama, A., Ogata, Y., Myler, P. J. and Stuart, K. (2008). Trypanosoma brucei Mitochondrial Ribosomes. Mol Cell Proteomics 7, 1286–1296.

