## Supplemental figures for "Nucleolar targeting in an early-branching eukaryote suggests a general physicochemical mechanism for ribosome protein sorting"

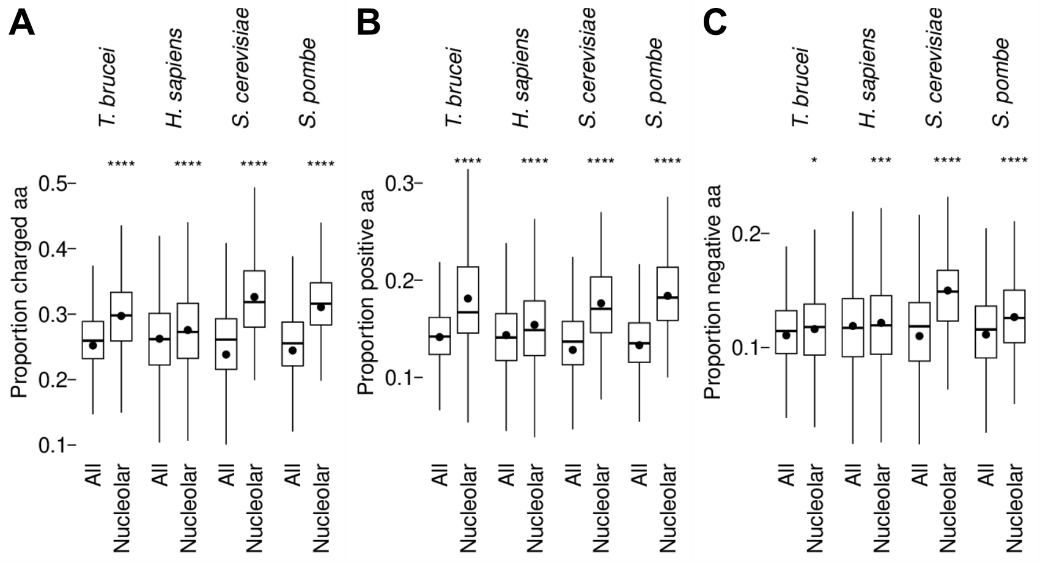

Figure S1. *T. brucei,* yeast and human nucleolar proteins tend to have many positively charged amino acids.

A. Proportion of charged (RHKDE) amino acids found in proteins annotated as nucleolar in *T. brucei*, in comparison an equivalent analysis of all proteins encoded in the genome from the *S. cerevisiae, S. pombe* and *H. sapiens* genome-wide protein localisation projects.

B. As for A, but for positively charged amino acids (RHK).

B. As for A, but for negatively charged amino acids (DE).

Statistical significance was assessed using the Wilcoxon signed-rank test (ns not significant, * p≤0.05, ** p≤0.01, *** p≤0.001, **** p≤0.0001).

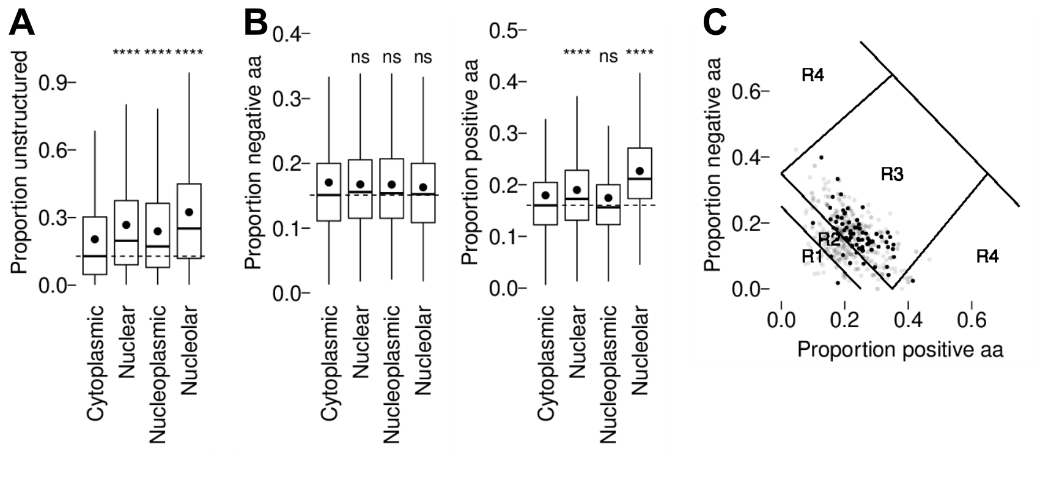

Figure S2. Unstructured domains in *T. brucei* nucleolar proteins.

A. Proportion of predicted intrinsically disordered (IUPRED) residues in cytoplasmic, nuclear, nucleoplasmic or nucleolar proteins, as classified by the cutoffs indicated in Fig. 2.

B. Proportion of positive (RHK) or negative (DE) charged amino acids found in the predicted unstructured domains of nuclear, nucleoplasmic or nucleolar proteins.

C. Diagram of states classification of predicted intrinsically disordered nucleolar protein domains. Strongly nucleolar (nucleolus/nucleus partition >1.2) are plotted in black. Specific regions are indicated: R1 globules, R2 globule and coil chimaeras, R3 polyampholytic coils or hairpins, R4 polyelectrolytic semi-flexible rods or coils.

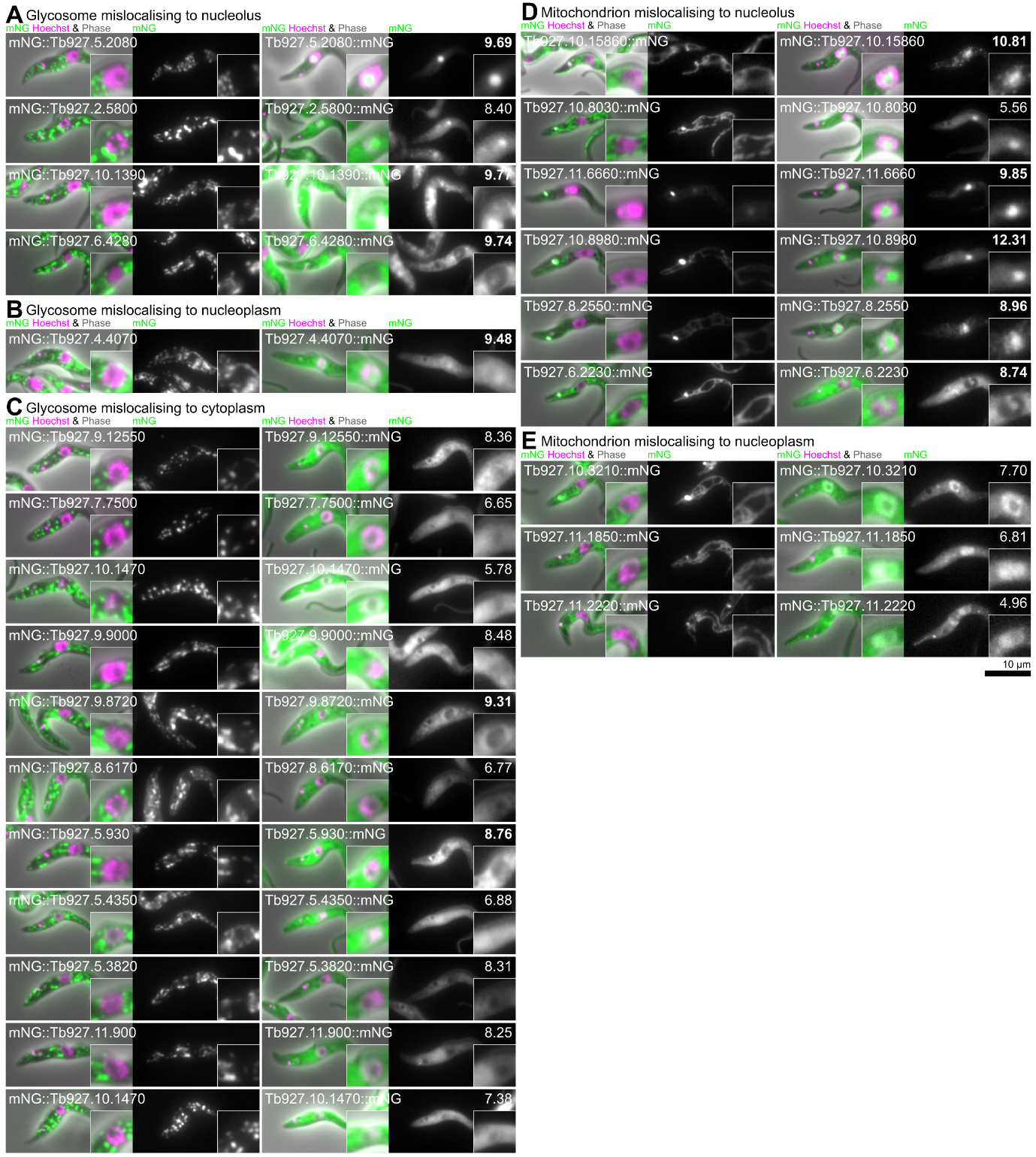

Figure S3. Basic proteins tend to mislocalise to the nucleolus when normal targeting sequences are disrupted.

A-C. Localisation of glycosomal proteins (with a C terminal glycosome/peroxisome targeting sequence) which mislocalise to the cytoplasm, nucleoplasm and/or nucleolus when tagged on the N terminus.

A. Glycosomal proteins which mislocalise to the nucleolus when tagged on the N terminus.

B. Glycosomal proteins which mislocalise to the nucleoplasm when tagged on the N terminus.

C. Glycosomal proteins which mislocalise to the cytoplasm when tagged on the N terminus.

D-E. Localisation of all mitochondrial proteins (with an N terminal mitochondrial targeting sequence) which mislocalise to the cytoplasm, nucleoplasm and/or nucleolus when tagged on the N terminus.

D. Mitochondrial proteins which mislocalise to the nucleolus when tagged on the N terminus.

E. Mitochondrial proteins which mislocalise to the nucleoplasm when tagged on the N terminus. No mitochondrial proteins mislocalise to the cytoplasm.

For each cell line, the mNG fusion protein pI is shown in the top right, in bold if >8.50.

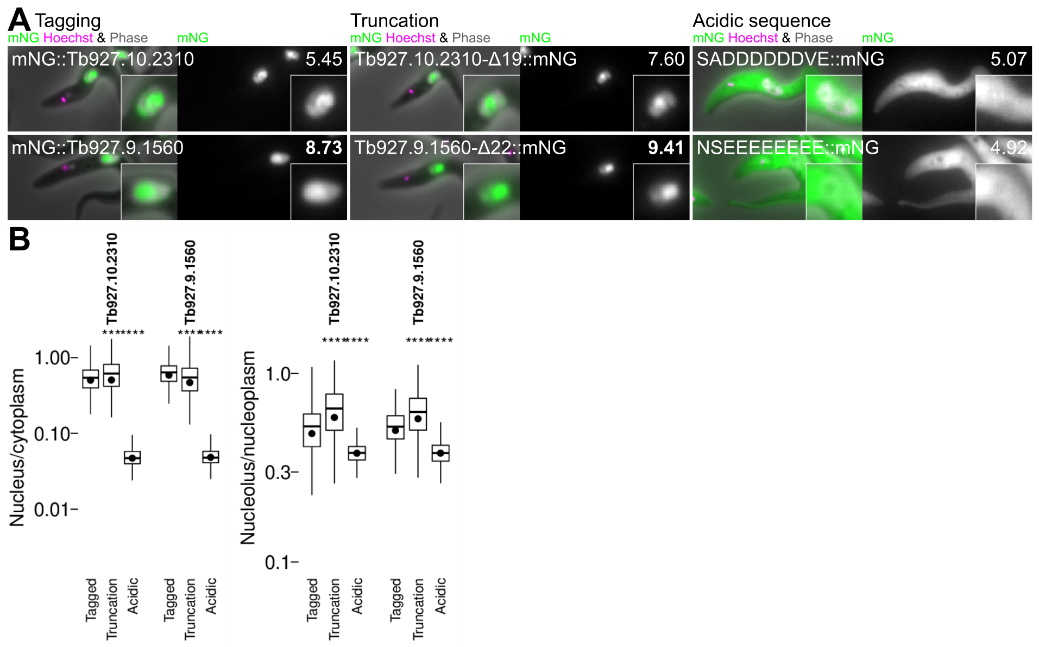

Figure S4. Acidic sequences in nuclear proteins are not required for nucleolar targeting.

A. Testing acidic runs for roles in nucleolar sequencing for 2 nuclear proteins with a single acidic run near the C terminus. Localisation of the protein by tagging at the endogenous locus, localisation following truncation to remove the C terminal acidic run and replacement with mNG and localisation of mNG fused to 10 amino acids of the acidic run. For each cell line, the number mNG fusion protein pI is shown in the top right, in bold if >8.50.

B. Plots of automated quantitation of the nucleus/cytoplasm and nucleolus/nucleoplasm mNG fluorescence signal partition from the cell lines in A.

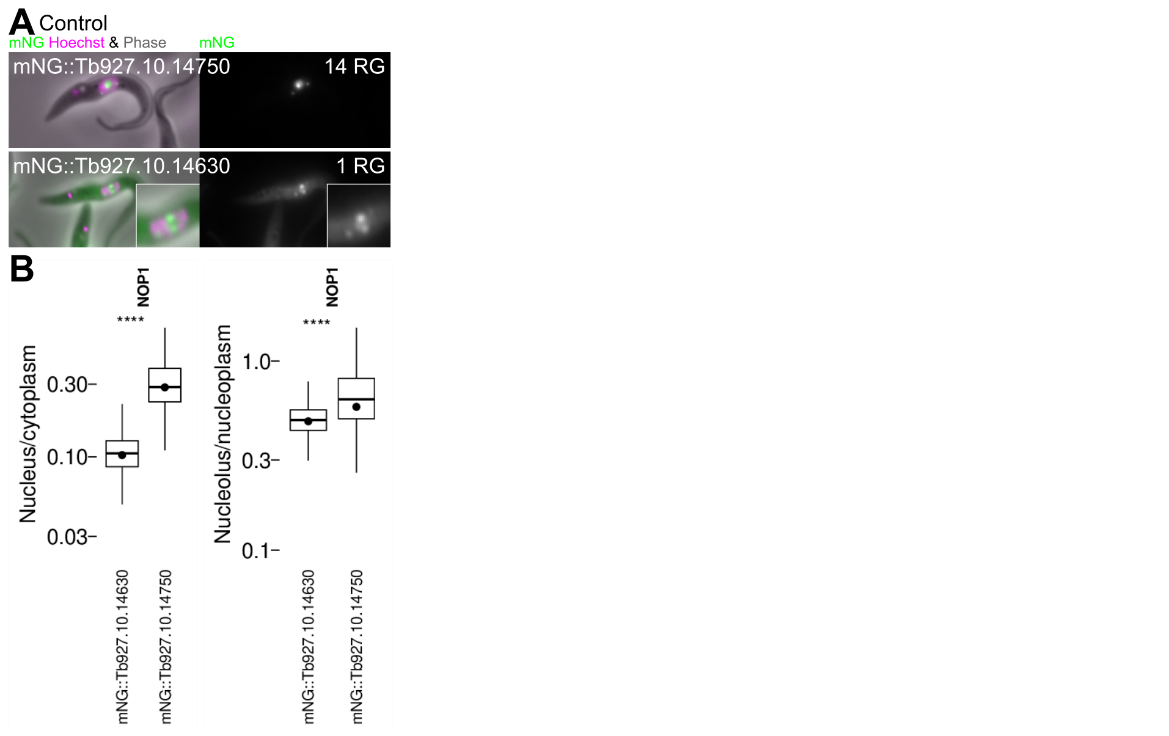

Figure S5. RGGs in nucleolar scaffolds are required for their partition to the nucleolus.

A. Localisation of two paralogs of *T. brucei* NOP1, one with 14 RGs in the sequence and one with an N terminal truncation leaving one RG.

B. Plots of automated quantitation of the nucleus/cytoplasm and nucleolus/nucleoplasm mNG fluorescence signal partition from the cell lines in A.
